# Characterizing white matter alterations in drug-naïve de novo Parkinson’s disease with diffusion MRI

**DOI:** 10.1101/2021.02.13.431081

**Authors:** Yiming Xiao, Terry M. Peters, Ali R. Khan

## Abstract

Parkinson’s disease (PD) is a progressive neurodegenerative disorder that is characterized by a range of motor and non-motor symptoms, often with the motor dysfunction initiated unilaterally. Knowledge regarding disease-related alterations in white matter pathways can effectively help improve the understanding of the disease and propose targeted treatment strategies. Microstructural imaging techniques, including diffusion tensor imaging (DTI), allows inspection of white matter integrity to study the pathogenesis of various neurological conditions. Previous voxel-based analyses with DTI measures, such as fractional anisotropy and mean diffusivity have uncovered changes in brain regions that are associated with PD, but the conclusions were inconsistent, partially due to small patient cohorts and the lack of consideration for clinical laterality onset, particularly in early PD. Fixel-based analysis (FBA) is a recent framework that offers tract-specific insights regarding white matter health, but very few FBA studies on PD exist. We present a study that reveals strengthened and weakened white matter integrity that is subject to symptom laterality in a large drug-naïve *de novo* PD cohort using complementary DTI and FBA measures. The findings suggest that the disease gives rise to both functional degeneration and the creation of compensatory networks in the early stage.

## 1. Introduction

Parkinson’s disease (PD) is a progressive neurodegenerative disorder characterized by the loss of dopaminergic neurons in the midbrain. Although the disease primarily affects the patient’s motor functions, it can also induce a range of non-motor symptoms, such as depression and sleep disturbance, as a result of modifications in brain connectivity (Ji *et al*., 2019). Precise knowledge regarding disease-related changes in white matter pathways can effectively help improve the understanding of the disease and its treatment strategy. Microstructural imaging techniques, including diffusion tensor imaging (DTI), allow inspection of white matter integrity to study the progression of neurological conditions.

In the past, a number of studies have investigated brain tissue changes in Parkinson’s disease using DTI techniques (Atkinson-Clement, Pinto, Eusebio, & Coulon, 2017; Zhang & Burock, 2020). Compared with morphometric analysis (Sarasso, Agosta, Piramide, & Filippi, 2020) that intends to characterize macroscopic anatomical changes, quantitative MRI measures have shown superior sensitivity in detecting tissue degeneration due to the disease. More specifically, voxel-based analyses (VBA) with DTI measures, such as fractional anisotropy (FA) and mean diffusivity (MD) have revealed PD-related tissue degeneration patterns, but unfortunately these findings were often inconsistent across studies, especially within white matter regions, partially due to smaller patient cohorts, sensitivity to heterogeneity in patient groups (e.g., variations in disease stages, symptoms, and treatment history), and MRI parameters (e.g., number of gradient directions in DTI sequences). In their meta-analysis of 39 studies, Atkinson-Clement *et al*. (2017) show that cellular degeneration measured as reduced FA and elevated MD exist within the substantia nigra, corpus callosum, and the cingulate and temporal cortices in PD patients.

Based on diffusion MRI, fixel-based analysis (FBA) (Raffelt *et al*., 2017) is a relatively recent method that offers tract-specific insights regarding white matter health. Instead of the more simplistic DTI model, FBA models fiber populations within one voxel (called a fixel) through functions of fiber orientation, and derives quantitative metrics in both macroscopic and microscopic scales to characterize white matter fibers, including fiber cross-section (FC), fiber density (FD), and the combined measure of fiber density and cross-section (FDC). Among these, FC reflects the macroscopic changes of the fiber bundles in terms of their cross-sectional areas; FD measures the density of fibers within a fiber bundle; and FDC evaluates the combined impact of FC and FD in white matter tracts. In contrast to conventional voxel-based analysis with DTI measures, which fails to handle fiber-crossing within voxels, FBA allows more interpretable results by identifying specific fiber tracts that are relevant to the designated study. Since its debut, this technique has demonstrated good specificity, sensitivity, and reliability in various applications, including natural aging (Choy *et al*., 2020), Alzheimer’s disease (Mito *et al*., 2018), stroke (Egorova *et al*., 2020), traumatic brain injury (Wallace *et al*., 2020), and mental disorders (Lyon *et al*., 2019).

To date, very few have attempted to characterize white matter alterations as a result of Parkinson’s disease using FBA. The existing studies (Li *et al*., 2020; Rau *et al*., 2019; Zarkali *et al*., 2020) primarily focus on early-stage patients, who have started drug treatment. More specifically, Rau *et al*. (2019) conducted a longitudinal analysis over a follow-up period of two years for 50 PD patients. Using 76 healthy controls to serve as the baseline, their results suggest an early degeneration in the splenium of the corpus callosum (CC), measured by the reduction of both FC and FDC in the region, as a typical alteration of PD. In addition to the CC, a decrease in FC or FDC was observed in several white matter tracts, including the cingulum, posterior thalamic radiation, corona radiata, cerebral peduncle, and internal capsule. Overall, fixel-based metrics were correlated with the worsening of a 39-item Parkinson’s disease questionnaire, Unified Parkinson’s Disease Rating Scale (UPDRS), and activity of daily living. Later, Li *et al*. (2020) compared fixel-based measures among 41 PD patients in the early stage (Hoehn & Yahr =1∼1.5), 57 PD patients in the middle stage (Hoehn & Yahr=2∼2.5), and 76 healthy controls. The comparison demonstrated significant differences in FD of the cerebrospinal tract (CST) and FC of the superior cerebral peduncle (SCP) among the three groups, with the mean FD values of the CST higher in the PD groups than healthy controls. In addition, they also found a correlation between reduced FD of CC and the deterioration of motor and psychiatric symptoms. Finally, Zarkali *et al*. (2020) investigated Parkinson hallucination and visual dysfunction with FBA, and discovered lower FC and FDC within the splenium of the CC, as well as lower FC in the left posterior thalamic radiata for patients with Parkinson hallucinations than those without the symptom.

It is known that motor symptoms of PD usually initiate unilaterally and then progress to both sides of the body, and evidence has linked the laterality of motor symptoms onset with the prognosis of the disease (Baumann, Held, Valko, Wienecke, & Waldvogel, 2014; Munhoz *et al*., 2013). For example, patients with the initial motor symptoms onset on the right-side are associated with faster disease progression (Baumann *et al*., 2014), and those with left-side onset were reported to have longer disease duration (Munhoz *et al*., 2013). Since the clinical laterality onset may impact the brain hemispheres differently (Scherfler *et al*., 2012; Shang *et al*., 2020), this factor should be fully considered in relevant neuroimage analyses of PD, especially when involving patients at the early stages of the disorder. Unfortunately, in a majority of current studies, the asymmetric development of PD is frequently overlooked. Besides the pitfall of limited patient cohorts in the analysis, the lack of consideration for laterality may also contribute to the inconsistency in the findings of disease-related tissue changes, thus preventing a clear depiction of pathogenesis of the disease. Therefore, an investigation based on a large PD cohort, with an emphasis on the clinical laterality onset should better elucidate the relationship between the development of PD and the associated alterations in white matter integrity. Furthermore, a comparison between FBA and VBA with DTI-measures based on the same cohort can help consolidate the findings with different analytical approaches. These are expected to be instrumental to further decode the disease mechanism and search for effective biomarkers for Parkinson’s disease.

In this study, we characterize the status of white matter health in a large cohort of drug-naïve *de novo* PD patients with respect to the laterality onset of motor symptoms using fixel-based analysis for the first time. From the public Parkinson’s Progression Markers Initiative (PPMI) database, 141 patients were compared against 62 sex- and age-matched healthy controls, as well as within the cohort to reveal strengthened and weakened white matter tracts, subject to the clinical laterality onset. In addition, we also compared the results from FBA to VBA with DTI metrics, which further supports the findings from FBA, while offering complementing insights.

## 2. Materials and Methods

### 2.1 Participants

For the study, 141 drug-naïve *de novo* PD patients (age = 61±9y, 51 females) having motor symptoms of unilateral dominance and 62 sex- and age-matched healthy subjects (age = 61±10y, 22 females) were obtained from the Parkinson’s Progression Markers Initiative database (www.ppmi-info.org/data), which is an ongoing multicenter study consisting of clinical evaluation, genetic information, bio-samples, and imaging data for the goal of identifying disease biomarkers. The up-to-date information on the study can be found at www.ppmi-info.org. At each participating PPMI site, written informed consent was obtained from all participants in the study. For the investigation, we included subjects who have T1w, T2w, and DTI images with the minimum age of 30 years from the database, and the patient data were collected from the baseline visits (N=152). Any subjects (N=8) with poor image quality by visual inspection were removed from the collection. All included PD patients have received clinical assessments, including the UPDRS for their symptoms and are at Hoehn and Yahr (H&Y) stage I or II without prior pharmaceutical treatment. Among these patients, 62 have motor symptoms more dominant on the left side and 79 on the right side. Patients affected by the motor dysfunction equally on both sides (N=3) were excluded from the study. For easy annotation, we will refer the PD patients with left-dominant symptoms as ***LPD*** and those with right-dominant symptoms as ***RPD***. From the clinical evaluations, the described study focuses on the UPDRS-I (non-motor experience of daily living), UPDRS-III (motor function examination), total UPDRS (the composite of motor and non-motor symptom evaluations), the Geriatric Depression Scale (GDS), and REM Sleep Behavior Disorder Screening Questionnaire Score (RBDSQ). They cover motor and non-motor symptoms of PD that are of particular interest in the early stage of the disease. The detailed demographic information of the subjects and the symptom evaluations are detailed in *Table 1*. The same information with respect to the dominant side of motor symptoms is listed in *Table 2*. Statistical tests were performed in each table to assess the differences in the demographic and clinical information between groups.

**Table 1.** Demographics of Parkinson’s disease and healthy subjects. Group differences were evaluated using statistical tests (a=Kruskal-Wallis test, b=Chi-square test, and c=Wilcoxon rank sum test), and *p-values* lower than 0.05 were considered significant (annotated with *).

|  | PD (n=141) | HC (n=62) | p-value |
| --- | --- | --- | --- |
| Age | 61.7±8.9 | 61.4±9.8 | 0.929<sup>a</sup> |
| Sex | 51 F & 90 M | 22 F & 40 M | 0.925<sup>b</sup> |
| UPDRS I | 5.1±3.8 | 2.4±3.3 | <1.0 x10<sup>-7</sup> <sup>c</sup> * |
| UPDRS III | 20.8±9.0 | 0.6±1.3 | <1.0 x10<sup>-29</sup> <sup>c</sup> * |
| Total UPDRS | 31.4±13.3 | 3.3±3.8 | <1.0 x10<sup>-28</sup> <sup>c</sup> * |
| GDS | 2.3±2.5 | 1.5±2.8 | 0.0014<sup>c</sup> * |
| RBDSQ | 3.9±2.5 | 2.9±2.3 | 0.0049<sup>c</sup> * |

**Table 2.** Demographics of PD patients with left dominant and right dominant symptoms. Group differences between left and right dominant PD groups were evaluated using statistical tests (a=Kruskal-Wallis test, b=Chi-square test, and c=Wilcoxon rank sum test), and *p-values* lower than 0.05 were considered significant (annotated with *).

|  | Left dominant<br>(n=62) | Right dominant<br>(n=79) | p-value |
| --- | --- | --- | --- |
| Age | 59.8±8.8 | 63.2±8.8 | 0.024<sup>a</sup>* |
| Sex | 28 F & 34 M | 23 F & 56 M | 0.049<sup>b</sup>* |
| UPDRS I | 5.2±3.5 | 5.1±4.0 | 0.702<sup>c</sup> |
| UPDRS III | 22.2±9.1 | 19.7±8.8 | 0.114<sup>c</sup> |
| Total UPDRS | 32.4±12.3 | 30.6±14.0 | 0.349<sup>c</sup> |
| GDS | 2.6±2.6 | 2.1±2.5 | 0.156<sup>c</sup> |
| RBDSQ | 3.8±2.1 | 4.0±2.7 | 0.945<sup>c</sup> |

### 2.1 Imaging protocols and pre-processing

For each subject included in the study, we acquired full brain T1w and T2w structural MRI volumes and diffusion MRI scans from the PPMI database. The T1w images were obtained using a magnetization-prepared rapid acquisition gradient echo (MPRAGE) protocol (TE=2.98ms, TR=2300ms, flip angle=9°, and resolution=1×1×1mm^3^). The T2w MRIs have an in-plane resolution of 1×1mm^2^ and slice thickness of 2∼3 mm. The diffusion-weight imaging (DWI) data were acquired using the following parameters: TE=88ms, TR=900ms, 64 gradient directions, b-values=1000s/mm^2^, and resolution=2×2×2mm^3^. For the DWI sequence, one B_0_ volume was acquired for each scan. The detailed MRI protocols can be found on the PPMI website (http://www.ppmi-info.org/study-design/research-documents-and-sops/).

Both T1w and T2w structural images were pre-processed with non-local mean image denoising (Manjon, Coupe, Marti-Bonmati, Collins, & Robles, 2010), non-uniformity correction using the N4 algorithm (Tustison *et al*., 2010), and intensity standardization. The brain mask was computed from T1w MRI using BeAST (Eskildsen *et al*., 2012), and T2w volumes were rigidly registered to the corresponding T1w images and resampled to 1×1×1mm^3^ resolution. DWI scans were processed with denoising (Veraart *et al*., 2016), unringing, eddy current correction (Andersson & Sotiropoulos, 2016), motion and distortion correction, and finally bias field correction (Tustison *et al*., 2010). More specifically, susceptibility-related distortion in DWI scans was corrected by first obtaining the deformation fields from nonlinear registration between B_0_ and T2w images in the phase encoding direction of DWI, and then applying the resulting deformation to the entire DWI series. The rest of the DWI pre-processing was performed using the MRtrix3 package (Tournier *et al*., 2019).

### 2.3 Fixel-based analysis

We use the single-shell 3-tissue constrained spherical deconvolution (SS3T-CSD) algorithm (Dhollander, Raffelt, & Connelly, 2016) implemented in MRtrix3 to conduct fixel-based analysis for our investigation. The algorithm can resolve white matter fiber orientation distributions (FODs) from single-shell diffusion MRI data while accounting for grey matter and cerebrospinal fluid (CSF) compartments. With the pre-processed DWI volumes, we obtained the FODs for each subject. In total, we conducted the following four groups of studies to characterize the white matter integrity using FBA, with the consideration of clinical laterality onset.

#### FBA Study 1

The FOD images from the LPD patients were left-right flipped while those from the rest of the patients and heathy controls were kept the same. This way, the clinical laterality onset of the patient population is kept on the right side of the body, and fixel-based metrics were compared between the PD and healthy groups.

#### FBA Study 2

With the clinical laterality onset aligned to the right, we took the data from all PD patients in *FBA Study 1*, and correlated the fixel-based metrics with the clinical scores UDPRS-I, UPDRS-III, total UPDRS, GDS, and RBDSQ.

#### FBA Study 3

The left- and right-dominant patients were separated into two groups. The fixel-based metrics of each group were compared against those of the healthy controls.

#### FBA Study 4

The left- and right-dominant patients were separated into two groups. Within each group, clinical evaluations, including UDPRS-I, UPDRS-III, total UPDRS, GDS, and RBDSQ, were correlated with the fixel-based metrics.

For each study, a study-specific template was created from all relevant subjects by using group-wise diffeomorphic nonlinear registration based on FODs at 1.3×1.3×1.3mm^3^ resolution. Then, the FOD image of each subject in the respective study was nonlinearly registered to the template, and the deformation fields were saved. A whole-brain tractogram with 2 million streamlines was generated by first obtaining an initial tractogram with 20 million streamlines using the probabilistic tractography algorithm, iFOD2 (Dhollander, Calamante, & Connelly, 2010) on the population-averaged FOD template and then filtering the result with the spherical deconvolution informed filtering of tractograms (SIFT) algorithm (Smith, Tournier, Calamante, & Connelly, 2013), to reduce false positive streamlines. In the template space, fixel-based measures of FD and FC were computed within each voxel for all subjects of interest, and their combined measure, FDC, was obtained by multiplying FC and FD. Note that for statistical tests, the logarithm of FC is used instead of the raw value. The generalized linear model (GLM) was used to test the hypotheses of each study, and the connectivity-based fixel enhancement (CFE) (Raffelt *et al*., 2015) was employed to identify fixel metrics that are associated with the factors to be tested. For all studies, the effects of sex and age were modelled as covariates and controlled for all the tests. Here, family-wise error (FWE) corrected p-values were computed for each fixel based on 10000 random shuffles to account for multiple comparisons, and the correct p-value < 0.05 is used to determine the statistical significance. Results with fewer than 10 fixels that showed statistical significance were not considered meaningful and thus not reported. The Johns Hopkins University (JHU) white matter tract atlas (Mori *et al*., 2008) was used to help guide the identification of relevant white matter tracts.

### 2.4 Voxel-based analysis with DTI metrics

To complement the results of fixel-based analysis with more commonly used measures of white matter integrity, we performed voxel-based analyses with fractional anisotropy (FA) and mean diffusivity (MD) for the same population. After DWI preprocessing, we calculated FA and MD maps for each participant using FMRIB’s Diffusion Toolbox (FDT, part of FSL). Each individual participant’s FA and MD maps were then transformed to the respective template space, using the corresponding warps generated from FBA in Section 2.3. We then performed voxel-based analysis in the template space using the GLM. The JHU white matter tract atlas (Mori *et al*., 2008) was used to constrain the region of interest (ROI) for the analysis and to help guide the identification of relevant white matter tracts. Similar to FBA, non-parametric permutation testing and FWE corrected p-values were used to reveal significant results (corrected p-value<0.05). The effects of sex and age were modelled as covariates and controlled for all the studies.

For voxel-based analysis, we also conducted four groups of studies, following the same strategies for FBA in Section 2.3. However, instead of testing fixel-based metrics, FA and MD values were analyzed instead. With the correspondence to the respective FBA studies, we denote the sub-studies for DTI metrics as ***VBA Study 1∼4***.

## 3. Results

### 3.1 Fixel-based analysis

With ***FBA Study 1***, we compare the fixel-based measures between PD patients and healthy controls after aligning the symptom dominant side to the right side of the body. The results are demonstrated in Fig.1, with both increase and decrease in fixel-based metrics in PD brains. More specifically, we have observed increased FC and FDC in a number of white matter tracts in the right hemisphere, including the posterior limb of internal capsule, cingulum, corticospinal tract, and superior cerebellar peduncles. In the left hemisphere, there was an increase in FDC in the anterior limb of internal capsule. Finally, the left cingulum exhibited lower FD in PD patients. The results of ***FBA Study 2*** are shown in Fig. 2. With the symptom dominant side aligned, we observed that the GDS score was inversely correlated with FD in the left stria terminalis and FDC in superior cerebellar peduncle.

**Figure 1.**
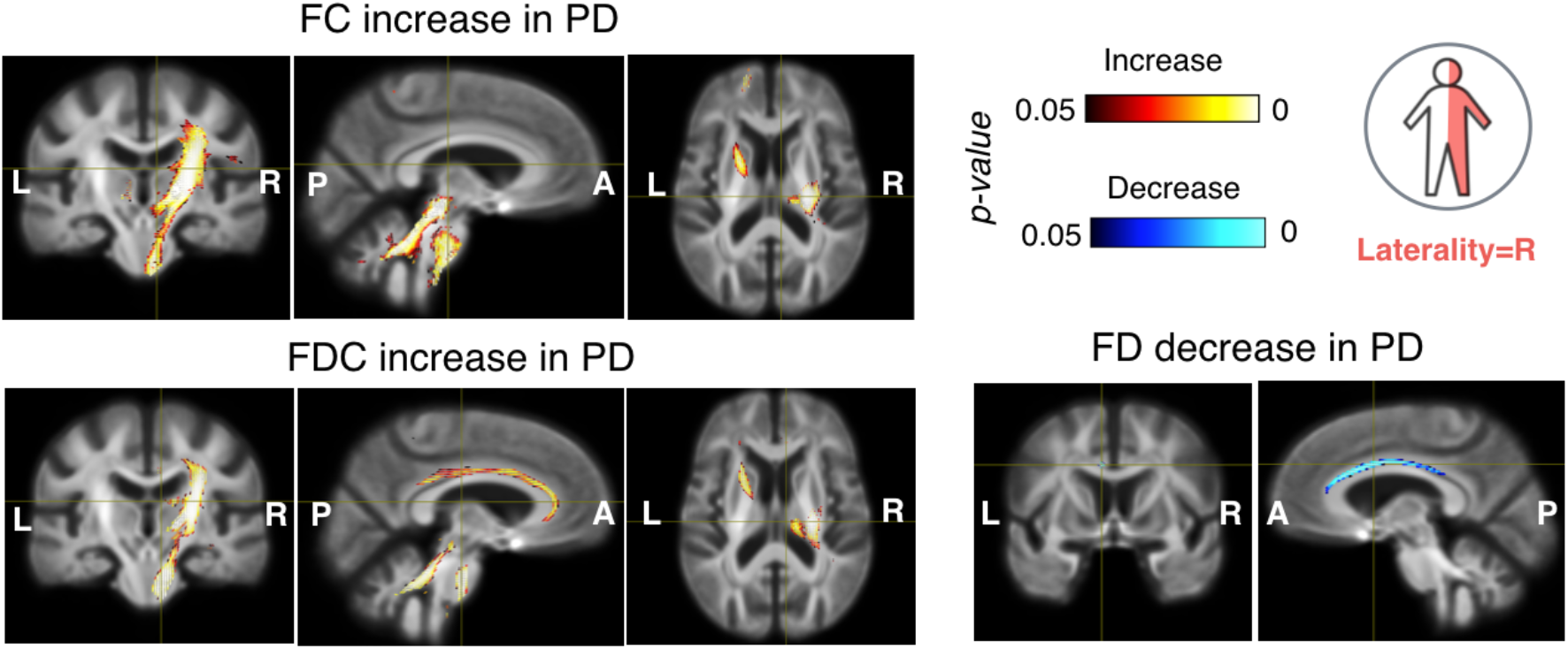
Comparison of fixel-based metrics (FC, FD, and FDC) between PD and healthy groups after aligning laterality of symptom onset (*FBA Study-1*). Significant fiber tracts of fixel-based metrics are overlaid on the group-averaged brain template. In each image group, the cross-hair points at the same location.

**Figure 2.**
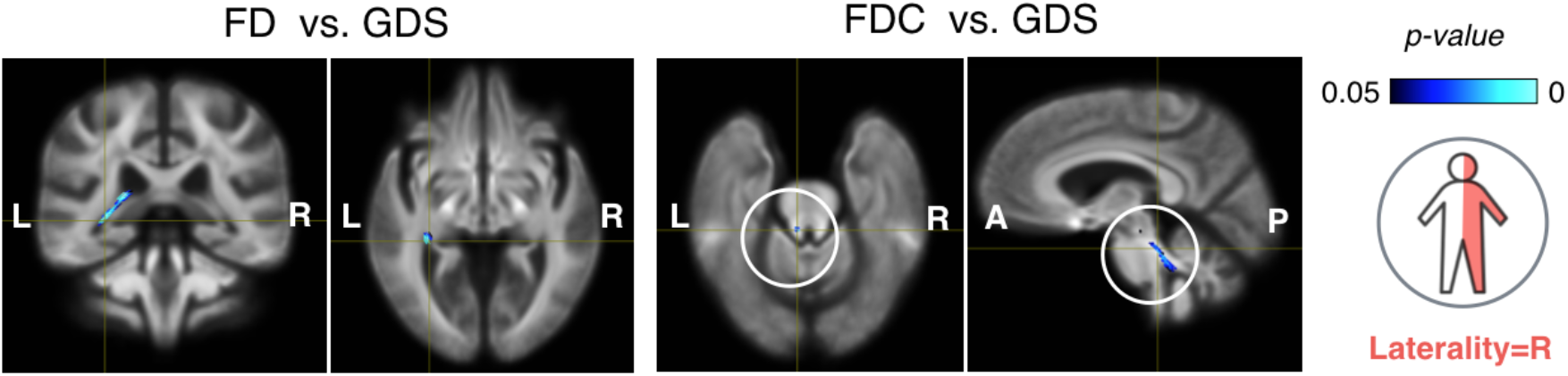
Correlation between fixel-based metrics and clinical evaluations in PD population after aligning laterality of symptom onset (*FBA Study-2*). Significant fiber tracts of fixel-based metrics are overlaid on the group-averaged brain template.

In ***FBA Study 3*** and ***FBA Study 4***, the PD cohort was divided into the left- and right-dominant groups. When investigating the differences between each PD sub-cohort and healthy controls in ***FBA Study 3***, higher FC in the right superior cerebellar peduncle and the cerebellum was observed in RPD (see Fig. 3) while no statistically significant results were detected for LPD vs. healthy subjects. When inspecting the association with clinical scores for the LPD sub-cohort (see Fig. 4), we found that there was a negative correlation between FC in the right dentatothalamic tract and the RBDSQ evaluation. On the other hand, for RPD (see Fig. 5), higher GDS was associated with decreased FD in the left stria terminalis and elevated FC in the splenium of the corpus callosum and fornix.

**Figure 3.**
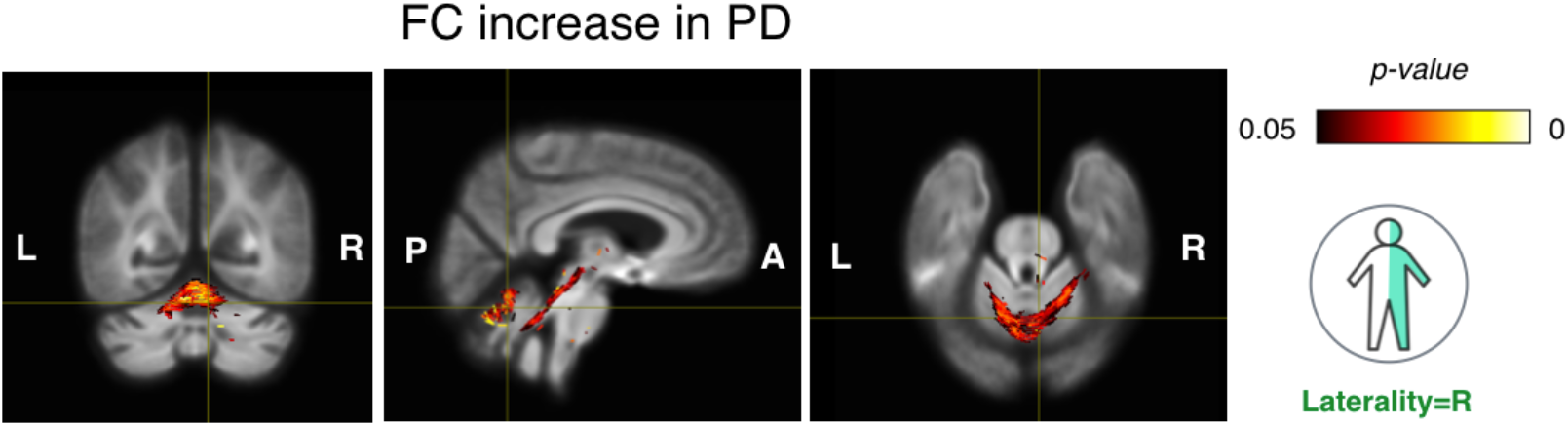
Comparison of fixel-based metrics between right-dominant PD and healthy groups after (*FBA Study-3*). Significant fiber tracts of fixel-based metrics are overlaid on the group-averaged brain template.

**Figure 4.**
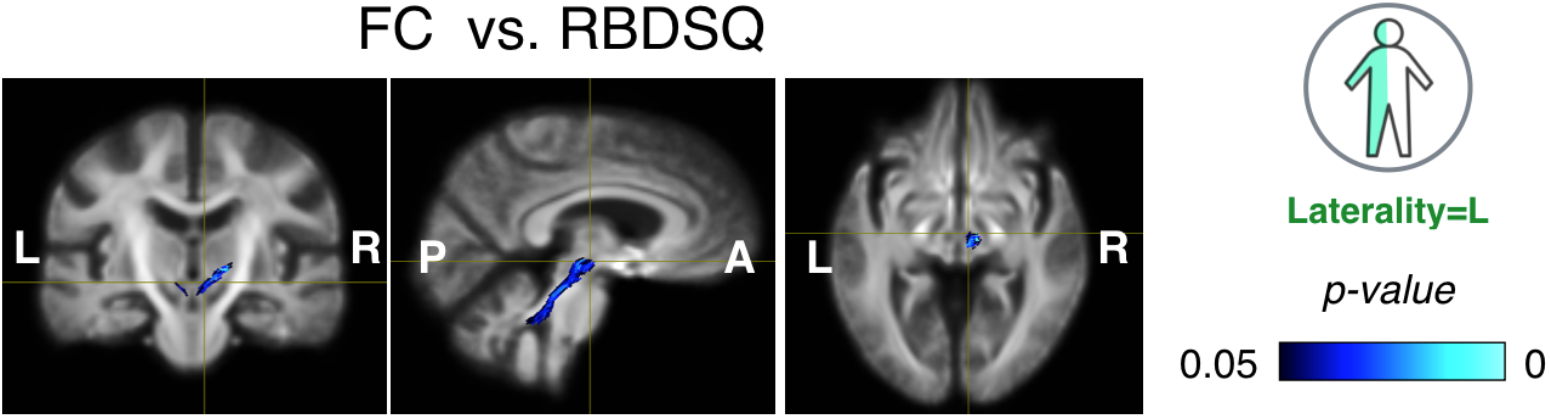
Correlation between fixel-based metrics and clinical evaluations in left-dominated PD cohorts (*FBA Study 4*). Significant fiber tracts of fixel-based metrics are overlaid on the group-averaged brain template.

**Figure 5.**
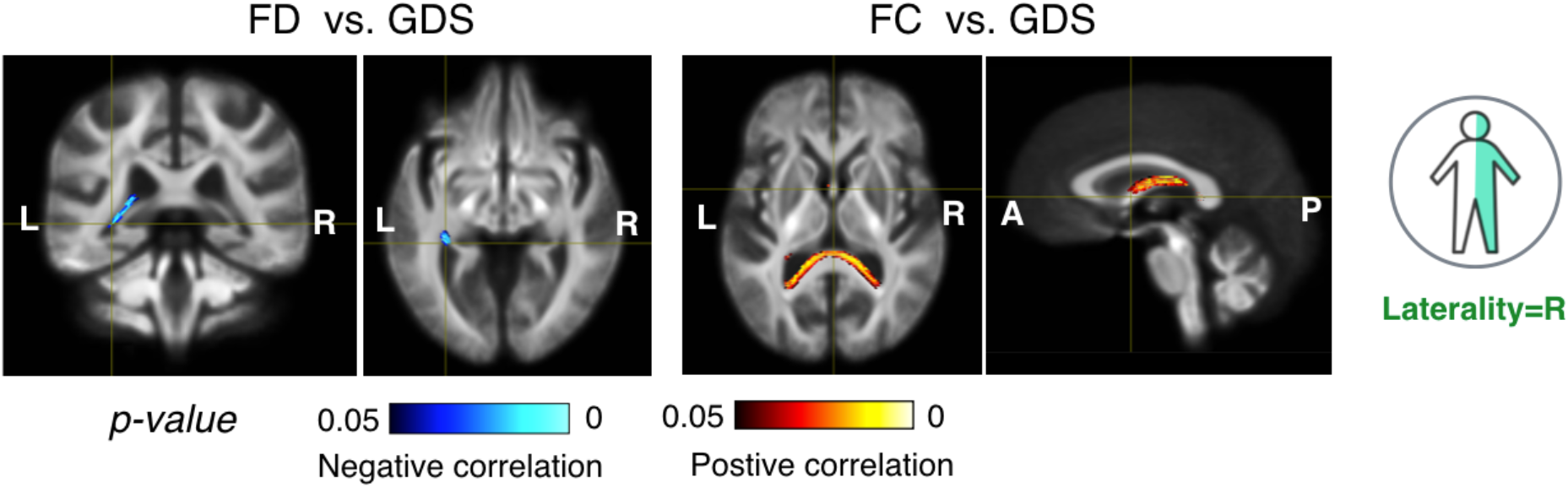
Correlation between fixel-based metrics and clinical evaluations in right-dominated PD cohorts (*FBA Study 4*). Significant fiber tracts of fixel-based metrics are overlaid on the group-averaged brain template.

### 3.2 Voxel-based analysis

We evaluated FA and MD in PD patients against those of healthy controls in ***VBA Study 1***, after aligning the symptom dominant side to the right-side of the body. The results are shown in Fig. 6. Compared with healthy subjects, PD patients had a widespread increase in FA primarily within the right hemisphere, which was ipsilateral to the dominant side of motor symptoms. The white matter tracts with FA increase included the right superior longitudinal fasciculus (SLF), posterior limb of internal capsule, cerebellar peduncle, cingulum, superior corona radiata, external capsule, and middle cerebral peduncle. In addition, there was also an increase of FA in the left anterior corona radiata, and reduction of MD in the patients occurred in the right internal capsule and superior corona radiata. On the other hand, MD increase in PD can be found in small regions within the left posterior limb of internal capsule and external capsule. With ***VBA Study 2*** (see Fig. 7), white matter fiber deterioration in terms of higher MD in the left external capsule, superior corona radiata, and SLF was linked to worsening of UPDRS-III scores. Furthermore, the left external capsule had increased MD that were correlated with the total UPDRS assessment.

**Figure 6.**
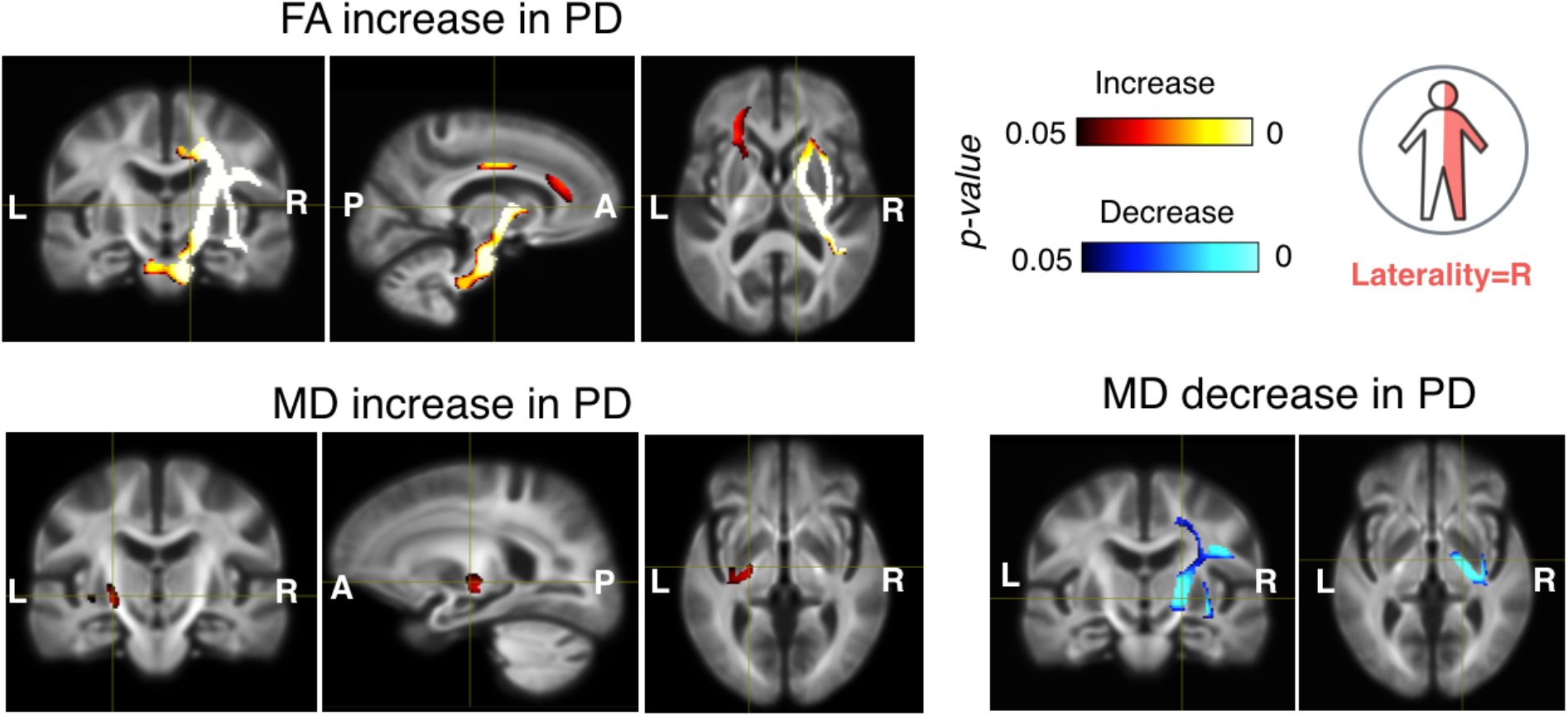
Comparison of fractional anisotropy (FA) and mean diffusivity (MD) between PD and healthy groups after aligning laterality of symptom onset (*VBA Study-1*). Significant clusters of DTI metrics are overlaid on the group-averaged brain template. In each image group, the cross-hair points at the same location.

**Figure 7.**
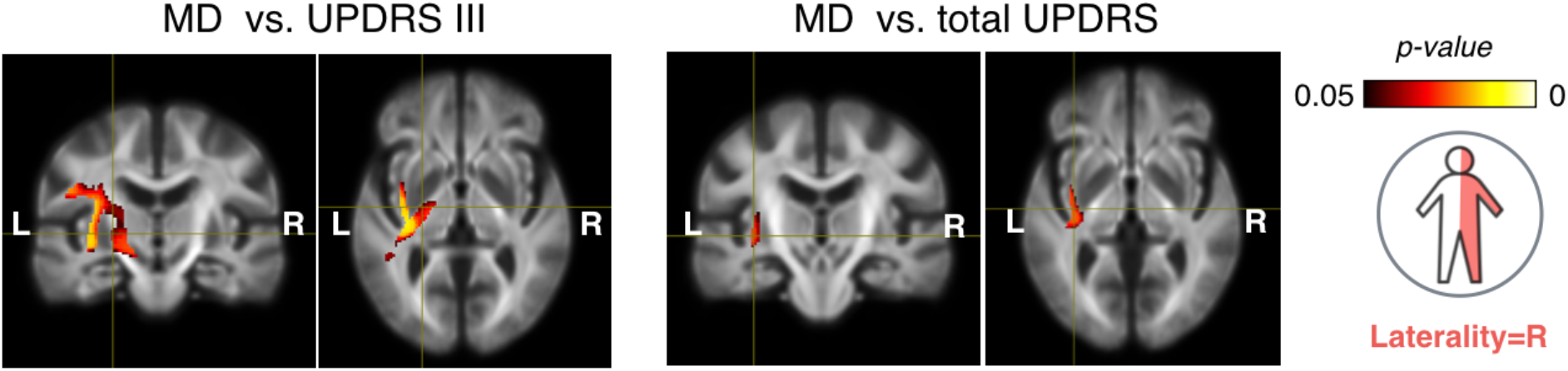
Positive correlation between DTI metrics (FA & MD) and clinical evaluations in PD population after aligning laterality of symptom onset (*VBA Study 2*). Significant regions are overlaid on the group-averaged brain template.

When comparing left- and right-dominant PD groups with the healthy controls (***VBA Study 3***), as demonstrated in Fig. 8, we found increased FA in the left posterior thalamic radiata and superior corona radiata in the LPD cohort while no differences were detected between the RPD group and healthy controls. With ***VBA Study 4***, several trends were observed in Fig. 9. In the LPD sub-cohort, the UPDRS-III score was associated with higher MD values in the left internal capsule, superior corona radiata, and external capsule. For RPD, MD values in the retrolenticular part of the internal capsule in the left hemisphere are positively correlated with GDS.

**Figure 8.**
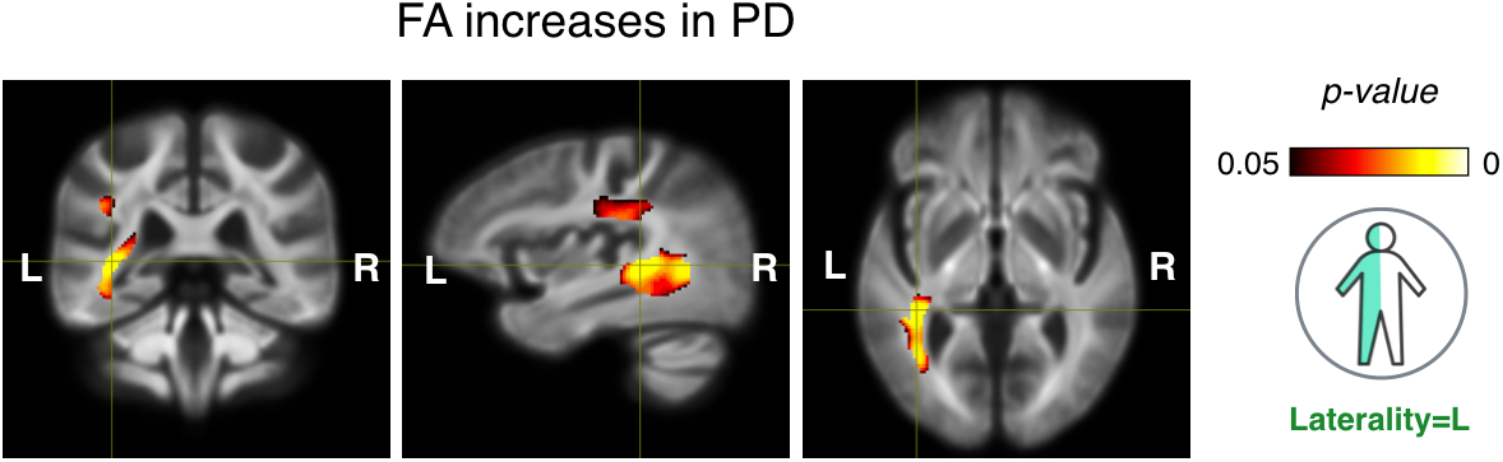
Comparison of FA values between PD and healthy groups in the left-dominant PD cohort (*VBA Study 3*). Significant regions are overlaid on the group-averaged brain template.

**Figure 9.**
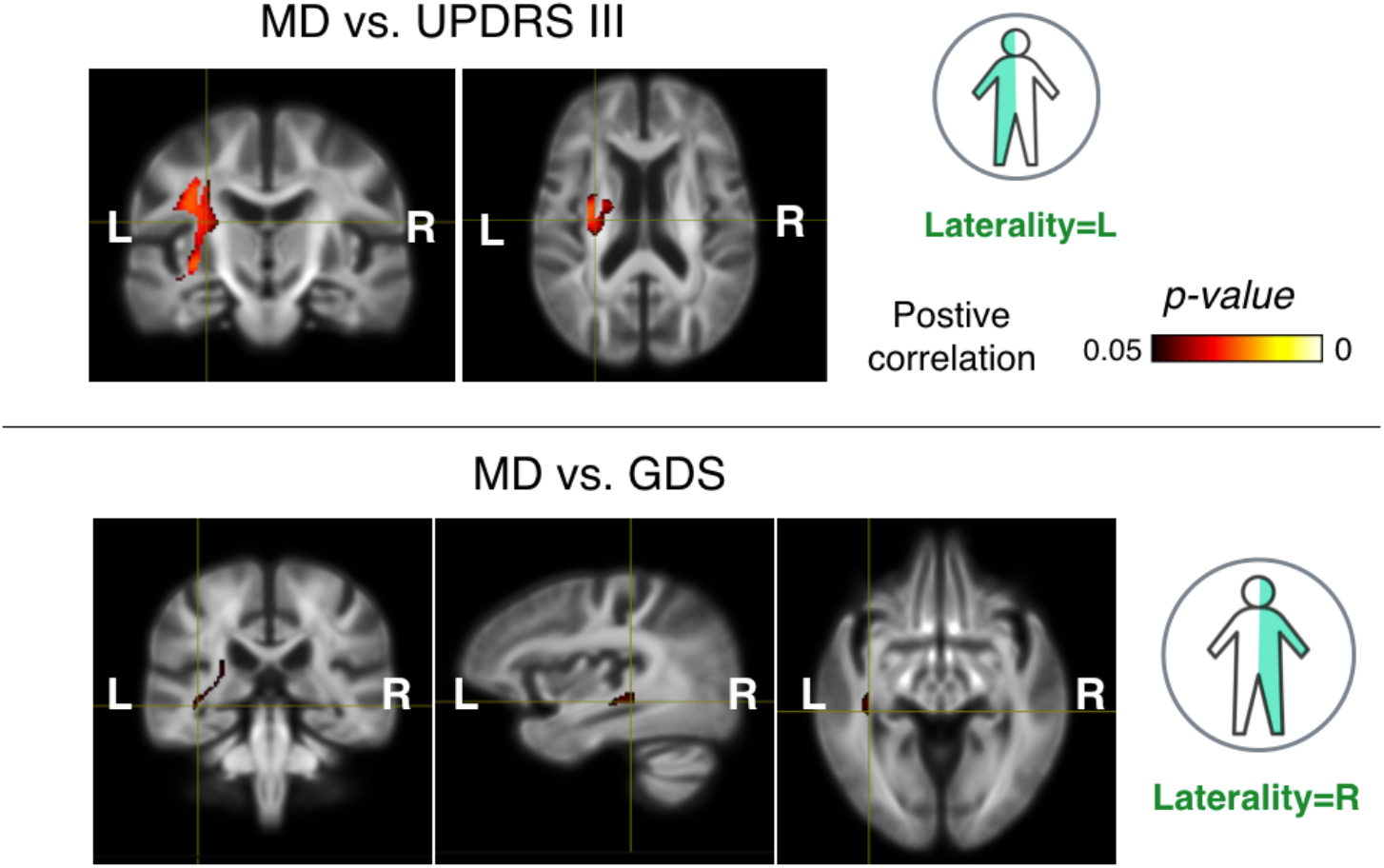
Correlation between DTI metrics (FA & MD) and clinical evaluations in left- and right-dominated PD cohorts (*VBA Study 4*). Significant regions are overlaid on the group-averaged brain template.

## 4 Discussion

In ***FBA Study 1***, where scans of LPD patients were flipped in the left-right direction, higher FC and FDC were observed in the right hemisphere for PD patients compared to healthy subjects, primarily in white matter tracts that are associated with motor control. This suggests a contralateral compensation of the brain to the initial motor function deterioration in early PD. The same trend is replicated in the elevated FA and decreased MD in the same hemisphere from ***VBA Study 1***. These results confirmed the observation in a few previous studies (Li *et al*., 2020; Mole *et al*., 2016), where higher FD and FA in CST were detected in PD compared to HC. However, while these studies suggested bilateral changes, we observed primarily unilateral alterations, potentially due to the consideration of symptom laterality onset. On the other hand, degeneration of white matter was also shown with both fixel-based and DTI metrics in the hemisphere that is more affected by motor dysfunction. Interestingly, while FBA identified the cingulum, which is more involved in the limbic system, VBA revealed regions that are more relevant to motor functions. When inspecting the clinical symptoms within the PD group (***Study 2***), with the same strategy of aligning the clinical laterality onset, white matter degeneration in the more affected hemisphere was associated with depression (i.e., GDS) by FBA, and with UPDRS III (motor dysfunction) and total UPDRS by VBA.

To better elucidate the distinct trends in LPD and RPD cohorts, the analyses in ***Study 3*** and ***Study 4*** examined these sub-groups separately. In ***FBA Study 3*** for RPD patients, elevated FC was observed in the right SCP, which is consistent with ***FBA Study 1***, and the cerebellum. Previous studies (Sweet *et al*., 2014; Wu & Hallett, 2013) have revealed the role of cerebellum and the related neural pathways in motor dysfunctions of Parkinson’s disease. As the RPD group have less severe clinical symptoms on average than LPD, whether the observed trend is specific to RPD or the disease progression stage will require further investigation. In ***VBA Study 3*** with LPD patients, regions with strengthened white matter integrity were found in the left hemisphere in PD patients. This further validated the implication of compensatory neuroplasticity in ***Study1***, but the higher FA in the posterior thalamic radiata was unique to LPD. When associating white matter health with non-motor symptoms of sleep disturbance and depression in ***VBA and FBA Study 4***, the relevant tissue degeneration also primarily occurred in the hemisphere with respect to the clinical laterality onset. However, in contrast to the trend in ***VBA Study 2***, there was a positive correlation between the UPDRS-III scores and the MD values in the left hemisphere for LPD. While lacking statistical significance between LPD and RPD, the mean UPDRS III and total UPDRS were higher in LPD than RPD. The observed correlation may be a result from the deterioration of motor functions on the less dominant side as the disease progresses. In general, FBA and VBA both offer insights of disease-related white matter changes, but since FBA emphasizes tract-specific measures, and VBM with DTI metrics produce more blobby regions, the identified regions with statistical significance don’t necessarily overlap with each other. Compared with FBA, VBA with DTI metrics is more sensitive in identifying associated brain regions with respect to clinical symptoms, despite the drawbacks in the difficulty in resolving crossing fibers.

In common with the existing FBA studies on PD progression (Li *et al*., 2020; Rau *et al*., 2019; Zarkali *et al*., 2020), both of which used 3T data, we have identified the splenium of CC, CST, SCP, cingulum, and fornix as regions that are affected by the disease. However, within these tracts, the specific trends in fixel-based measures don’t always agree among previous studies (Li *et al*., 2020; Rau *et al*., 2019) and ours. For example, Rau *et al*. (2019) discovered that the UDPRS assessments, including UPDRS III and total UPDRS, are positively correlated with FC in the body of CC, but Li *et al*. (2020) found a negative association between FD values in the same structure and UPDRS III. In our cohorts, no significant results were revealed for UPDRS evaluations and fixel-based measures. Furthermore, Li *et al*. (2020) associated reduced FD in the CC with depression while our results showed higher FC in the splenium of CC with depression in RPD. Previously, Pelizzari *et al*. (2020) have also compared DTI metrics between PD patients with unilateral symptoms and healthy controls. Different from our results, they found widespread white matter integrity reduction in terms of higher MD in RPD but not in LPD. These differences may originate from several sources, such as variations in imaging protocols (e.g., multi-shell vs. single-shell), cohort size, and patient heterogeneity. Notably, these recent studies included patients at slightly more advanced disease stages (H&Y stages 1∼3) than our PD cohort (H&Y stages 1∼2). Another potential contributor to the difference is the influence of dopaminergic therapy. This factor may alter white matter properties in addition to the disease itself, but is often overlooked in most studies (Zhang & Burock, 2020), including those mentioned in this paragraph. As the influence of drug therapy is still poorly understood, investigation of tissue changes in drug-naïve PD can provide a baseline to clarify the relationship between brain plasticity and PD treatment.

The development and progression of Parkinson’s disease is complex and nonlinear (Li *et al*., 2020; Rau *et al*., 2019; Sarasso *et al*., 2020), often with the motor symptoms initiating unilaterally and progressing to both sides in more advanced stages. This is expected to induce differential patterns of tissue changes, and thus brain connectivity alterations between the left and right hemispheres. However, most PD-related investigations on micro- and macro-structural changes do not consider the laterality of the dominant symptoms, especially when many focus on early stages of PD. Such an approach may reduce the sensitivity for detecting disease-related brain regions, or reporting bilateral changes instead of potentially unilateral changes, possibly contributing to inconsistent conclusions in the literature. Our analyses were conducted with strategies that fully account for laterality. In the first strategy, the sides with more severe symptoms were mapped to the right for all subjects. This allows the entire PD population to be included, thus boosting the statistical power to detect relevant trends with a clinical DTI protocol. As a result, we were able to reveal laterality-dependent microstructural alterations in drug-naïve *de novo* PD patients, which were not demonstrated previously. In the second strategy, sub-groups of LPD and RPD patients were analyzed separately. This approach helped confirm the insights related to laterality of symptoms onset, and provided complementary tissue characterizations that are specific to each sub-group. To inspect our first approach of aligning the side of symptoms onset, in Fig. 10, we illustrate the t-value maps from testing the hypothesis that the FC values are higher in the PD group than the healthy controls in ***FBA Study 1*** *and* ***3***. In the figure, the coronal slice from Fig.1 demonstrating significant FC alteration in motor-function-related tracts is shown with the corresponding t-value maps from the two studies. Despite the fact that the LPD and RPD sub-cohorts didn’t show statistically significant trends in the same white matter tracts, we can still observe a similar trend of higher t-values in the relevant regions from the hemisphere that is less affected by PD, across the three maps. Note that for easy visual comparison, the t-value map for the LPD group is left-right mirrored, and different t-value windows are used between studies.

**Figure 10.**
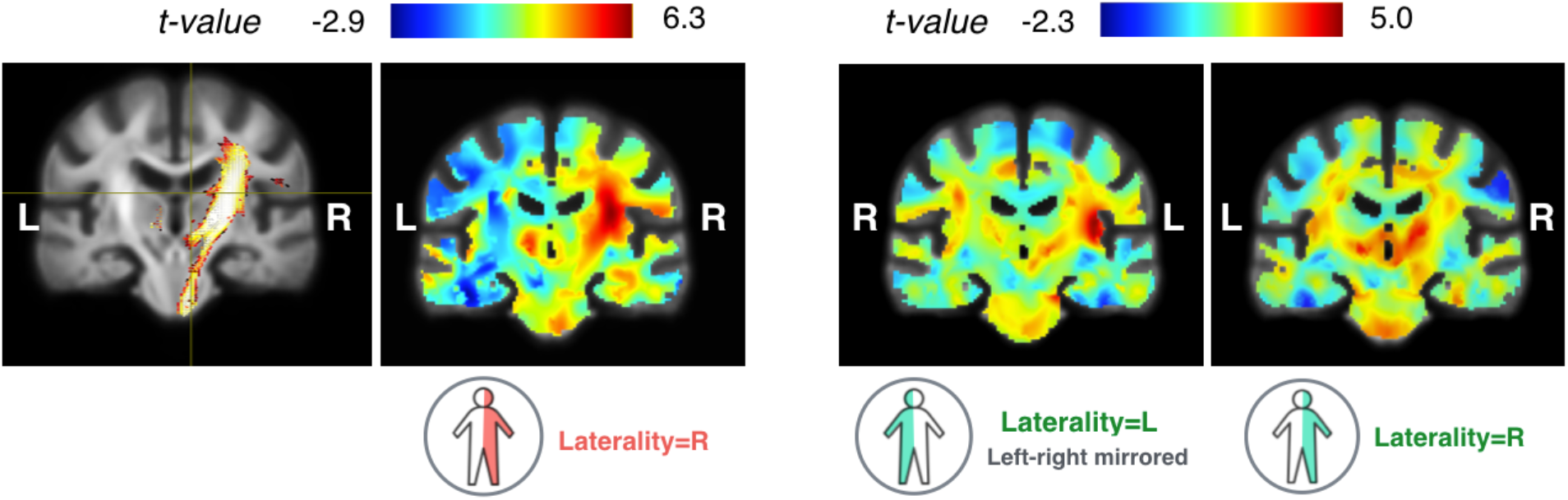
Demonstration of t-values from testing the hypothesis that the FC values are higher in PD cohorts than healthy controls in FBA *Study1 & 3*. From left to right: a coronal slice of the significant fixels from Fig.1, the corresponding t-value map from *Study1*, and the corresponding t-value maps from *Study3* with respect to LPD and RPD sub-cohorts. For easy visual comparison, different t-value windows are used between studies, and that of the LPD sub-cohort was left-right mirrored.

The described studies still have a number of limitations. First, the analyses employed clinical single-shell DTI protocols at 3T (64 gradient directions, b = 1000 s/mm^2^), and thus the data quality can be lower than those from multi-shell protocols (Rau *et al*., 2019; Zarkali *et al*., 2020), in terms of signal-to-noise ratio and angular contrast-to-noise ratio. For FBA, we employed the recent single-shell multi-tissue CSD algorithm to accommodate the scanning protocol, and showed that meaningful results can still be obtained with single-shell DTI scans. Second, as the DTI scans in the PPMI database do not have matching scans with opposing phase encoding directions, susceptibility-induced distortion was corrected through nonlinear registration between the corresponding B_0_ and T2w MRIs. Although this type of approach was used previously (Ardekani & Sinha, 2005), it may limit the accuracy of the analyses, particularly in regions with more severe distortions (e.g., orbital frontal and temporal lobes). Lastly, part of our study aligned the clinical laterality onset to the right side for analysis. In addition to the asymmetry introduced by the disease, inherent asymmetry of the brain may have also contributed to the analysis. To inspect the potential difference, LPD and RPD sub-groups were analyzed separately, and similar trend was found to the studies that align the dominant symptom side. However, further quantification of the influence from inherent brain asymmetry on PD is beyond the scope of the current investigation, and will be explored in detail in the future study.

## 5 Conclusion

In this study, we present the first investigation that characterizes white matter integrity with the consideration of symptom laterality in a large drug-naïve *de novo* PD cohort, using complementary DTI and FBA measures. In addition to revealing tissue alterations between PD and healthy subjects, we demonstrated that both motor and non-motor clinical assessments were correlated with different DTI and FBA measures within white matter pathways. The findings suggest that the disease gives rise to both functional degeneration and the creation of compensatory networks in the early stage, and the impacts are subject to the laterality of the motor symptoms. As the first FBA investigation with a drug naïve PD population, the insights are instrumental in understanding the progression of the disease and potentially inform the future exploration regarding the interplay between disease-related brain tissue changes and drug therapy. To depict a richer picture of the disease, future longitudinal studies will help fully characterize the trajectory of PD-related white matter changes as a potential biomarker for improved prognosis and treatments.

## Acknowledgements

The work is supported by the BrainsCAN and CIHR fellowships for Y. Xiao.

Yiming Xiao, Terry M. Peters, and Ali R. Khan have no conflict of interest.

PPMI—a public-private partnership—is funded by the Michael J. Fox Foundation for Parkinson’s Research and funding partners, including AbbVie, Avid, Biogen, Bristol-Myers Squibb, Covance, GE Healthcare, Genentech, GlaxoSmithKline, Lilly, Lundbeck, Merck, Meso Scale Discovery, Pfizer, Piramal, Roche, Sanofi Genzyme, Servier, Teva, and UCB.

The authors would like to thank Dr Thijs Dhollander for his insightful discussion on MRtrix software and fixel-based analysis.

